# Spatial generalization of peripherally encoded memories emerges before selection for report

**DOI:** 10.1101/2025.04.30.651449

**Authors:** G. Kandemir, C.N.L. Olivers

## Abstract

Visual working memory has been shown to transition from location-specific to spatially generalized representations. It remains unclear what the time course of this generalization is and to which information processing stages and signals it is linked – specifically whether it is tied to sensory or report stages. Here we used EEG to investigate spatial generalisation of peripherally encoded stimuli before and after an item was selected for report. Thirty healthy participants memorized two orientations presented serially at fifteen degrees eccentricity to the left and right of central fixation. One of the two items was subsequently cued as task relevant. During the delays, impulse signals were inserted to trace memoranda in potentially activity-reduced states. We observed that prior to the cue, patterns of activity transitioned from location-specific spatially generalized states from approximately 400-600 ms after onset in broadband voltage but not in oscillatory alpha signals. The picture was more mixed after the selection cue, with some evidence for locations-specific and spatially generalized codes in both alpha and broadband voltage. We conclude that the first stages of spatial generalization occur at sensory, pre-report levels of processing, while late stages are associated with selecting response-relevant spatial features.

## Introduction

Visual working memory (VWM) enables us to maintain visual information in a directly accessible state, readily available to guide behavior (Cowan, 2001). Neurophysiological studies have suggested that representations held in VWM spatially generalize, as evidenced by the observation of feature-specific activity patterns spreading beyond those tuned to the stimulus at its original location (Bae & Luck, 2018; Ester et al., 2009; Fukuda et al., 2016; Kandemir & Olivers, 2024; Kandemir et al., 2024a; see also Chambers et al., 2013; Jehee et al., 2011; Serences & Boynton, 2007; and Williams et al., 2008 for a similar spatial transfer effects in perceptual tasks rather than VWM). The observation of such generalized patterns is important, as they may inform us about the representational format in which such memories are being maintained, for example as sensory traces, report-related representations, or something in between (cf. Myers, et al., 2017; Olivers & Roelfsema, 2020; van Ede & Nobre, 2023).

Relatively little is known about during which processing stage(s) spatial generalization in VWM occurs. So far, studies addressing spatial generalization have either employed only a single to-be-remembered stimulus on each trial (typically an orientation; Bae & Luck, 2018; Ester et al., 2009; Fukuda et al., 2016; Kandemir & Olivers, 2024), or have only assessed generalization after observers were cued to select one of two memories (Kandemir et al, 2024a). This then also automatically makes the item relevant for report. It is possible that generalization then reflects the transition from a location-specific sensory representation to a representation that supports the task that participants need to conduct, as such a transition may involve certain levels of abstraction, and eventually conversion into response-supporting representations (cf. Bae & Luck, 2018, 2019; Bae & Chen, 2024; Cochrane & Green, 2023; Duan & Curtis, 2024; Henderson et al., 2022; Kandemir et al., 2024a; Kwak & Curtis, 2022; Mostert et al., 2018; Myers, et al., 2017; Olivers & Roelfsema, 2020; Panichello & Buschman, 2021; van Ede & Nobre, 2023). For example, spatial generalization may be expected when stimuli presented at different locations eventually map onto the same attentional process and/or the same response.

Here we sought to investigate whether spatial generalization is indeed tied to selection for report, or reflects a more universal mechanism that can already occur earlier. To this end we combined electroencephalography (EEG) with the retro-cue paradigm illustrated in Figure 1A. Participants memorized two serially presented orientations, each presented at 15° eccentricity on opposite sides, left and right from fixation, and each followed by a 1400 ms delay period. The patterns were thus clearly spatially and temporally separated. Only after the second delay, a cue indicated (with 100% validity) which of the two items was to be reported upon, followed by another delay. Finally, participants reported whether a probe stimulus was rotated clockwise or counterclockwise relative to the cued memorandum, and this direction was randomized. We were most interested in when spatial generalization would emerge. Importantly, prior to the cue, participants had no information on which orientation would be task relevant. Nor was there any information throughout the trial on what to respond until the probe was presented. Thus, if spatial generalization occurs only after the cue, then this would provide evidence consistent with the idea that selecting the item for report is crucial. In contrast, if such generalization already occurs prior to the cue, then selection for report is not, or not the only contributor to generalization.

**Figure 1.**
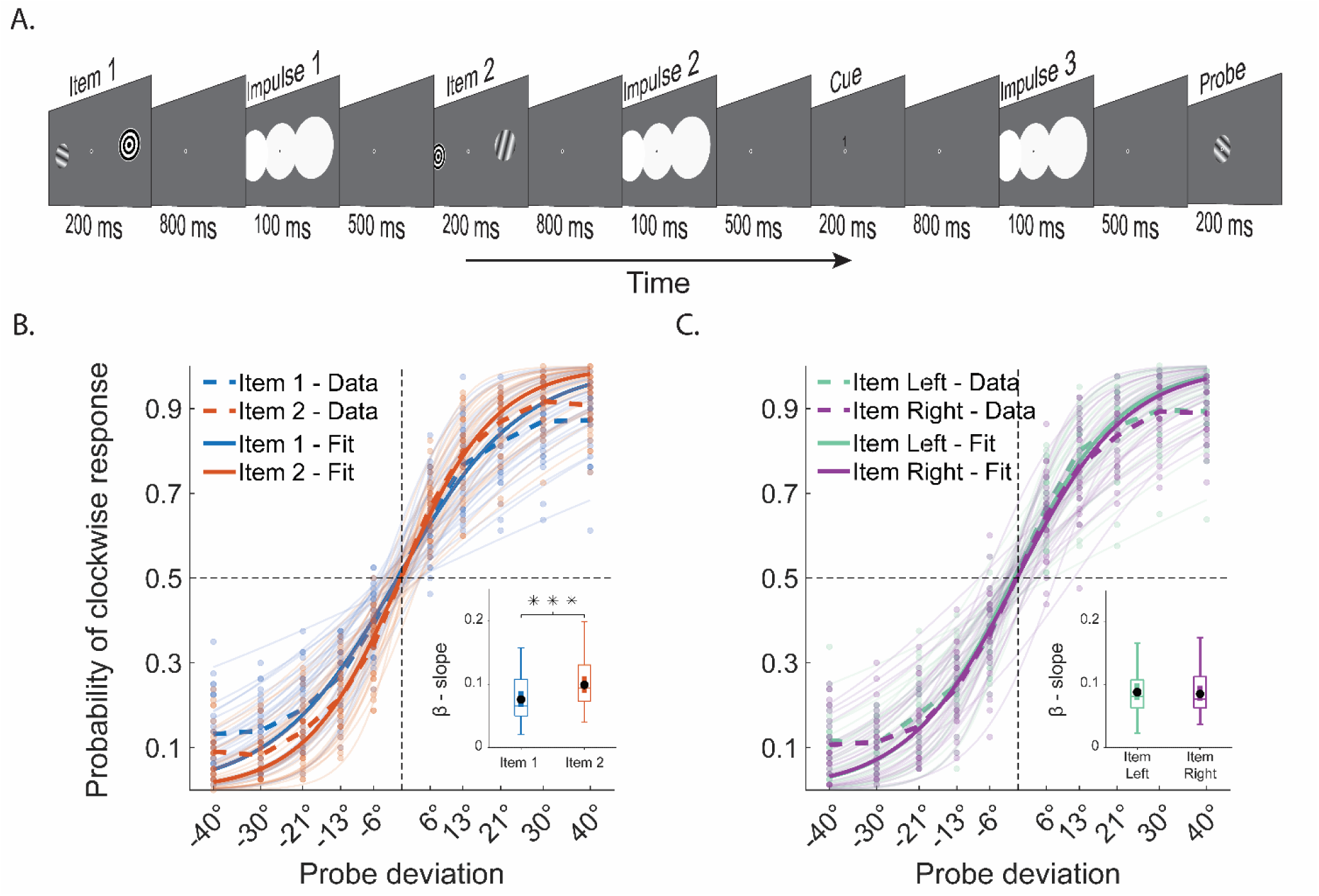
Illustration of a trial (A) performance split for item 1 and 2 (B), or split for left and right position (C), as a function of the true deviation of the probe from the target. Note. Demonstration of a single trial and task performance split for item location and order. **A.** In this example Item 1 is presented on the left, Item 2 on the right, and then Item 1 is retrospectively cued as task relevant (cues were “1”, or “2”, corresponding to the order of memory item presentation, and were always relevant). The report task required participants to compare the probe orientation to the cued item and indicate the relative direction (clock-wise or anti-clockwise). **B.** The probability of responding clockwise to the probe, split up for Item 1 (blue) and Item 2 (orange). **C.** The same but split up for items on the left (green) versus right (purple) side of the display Solid lines show the overall logistic fit for Item Left (green-solid) and Item Right (purple-solid) estimated using the average of individual parameters. A-B. Dashed lines who the data, solid lines the logistic fit (the average of the individual parameters). Faded dots and lines reflect the individual data and logistic fits. Boxplots compare slopes (β) across conditions where black dots mark mean slope and asterisks indicate significant difference (*** = *p* < 0.001).

Using a decoding approach, we focused on two types of signal, namely the *broadband voltage* (i.e. evoked potentials), and on the *oscillatory alpha power*, as these have been associated with different types of representation. Specifically, the broadband voltage been shown to be more consistently associated with the actual stimulus feature (here orientation), while alpha has been more consistently associated with either spatial abstractions – specifically directing spatial attention to part of the stimulus and/or preparing an action in a certain direction (Bae & Luck, 2018; 2019; Bae & Chen, 2024; Fahrenfort et al., 2017; Günseli et al., 2019; Hanslmayr, 2011; Kandemir & Olivers, 2024; Klimesch, 2012; Liu et al., 2022; van Moorselaar et al., 2018; Pavlov & Kotchoubey, 2022; de Vries et al., 2018). Given this distinction we may expect that if generalization emerges prior to the cue, when no information is available on which stimulus is response-relevant, it will primarily emerge in the broadband signal associated with sensory representations. After the cue, generalization is then expected to emerge more strongly in the alpha signal, when some shared attentional or response-related mapping can be created. Note here that a concrete response-related code is not a given in the current task, because it is not clear what the response will be until the probe stimulus is presented. Hence, any alpha-related representations may mainly reflect spatial attention-related activity.

Finally, given that activity patterns may subside during the delays, we also included three impulse stimuli (white flashes covering the two peripheral stimulus locations as well as central fixation), one in each delay period, in order to enhance our ability to trace memories in potentially activity-silent or -reduced states (Kandemir & Akyürek, 2023; Kandemir et al., 2024a; 2024b; Kandemir & Olivers, 2024; Stokes, 2015; Wolff et al., 2017; 2020a; 2020b; see Figure 1A).

## Methods

### Participants

The sample of participants consisted of thirty healthy young subjects (18 female, M_Age_ = 22.5, s.d._Age_ = 4.75), who achieved at least 65% accuracy in a shorter prescreening version of the experiment (160 trials) conducted prior to the main experiment. Participants were paid 13 euros/hr. The sample size was the same as earlier studies to acquire enough power to successfully discriminate orientations presented in the periphery (Kandemir & Olivers, 2024). Both prior to prescreening and the actual experiment, participants were informed about the purpose and the design of the study as well as the data maintenance practices verbally and in writing. A written consent form was signed by participants to declare their compliance with participation and data storage practices. Procedures were approved by the Scientific and Ethics Review Board of the Faculty of Behavioral and Movement Sciences at VU University Amsterdam (umbrella protocol number VCWE-2021-173).

### Stimuli and Apparatus

The design of the experiment and the duration of stimuli and delays were based on earlier studies using impulse driven EEG decoding (Wolff et al., 2017; Kandemir & Akyürek, 2023; Kandemir et al., 2024a; 2024b; Kandemir & Olivers, 2024). In each trial, two sinusoidal gratings were presented serially as memory items, each with a diameter of 5 degrees of visual angle (dva). The phase of the gratings varied randomly between 1 and 180 degrees for each trial, while the spatial frequency and contrast remained constant at 1. The orientations of the gratings that would then be cued were pseudo-randomly sampled from a set of eight discrete orientations, ranging from 11.25° to 168.75°, with an angular distance of 22.5° between them. The distractor orientation was randomly sampled from the set of eight with equal probability, in each block. The memory items were presented at an eccentricity of 15 dva to the left or right of the center of the screen (center-to-center distance). The location opposite the memory item was occupied by a bull’s-eye symbol consisting of concentric black and white circles (5 dva in diameter), serving as a placeholder.

The cue was either the number 1 or 2, displayed in black 12-point Arial font, 0.25 dva above the fixation dot at the center, indicating the serial position of the cued memory item. The probe was a sinusoidal grating presented at the center of the screen with a random phase ( 1 to 180 degrees) and fixed contrast and spatial frequency (both set to 1). The diameter of the grating was 5 dva. The orientation of the probe always differed from the cued memory item in increments of +/− 6 °, 13 °, 21 °, 30 ° or 39 °.

Three impulse stimuli presented at different timepoints of the experiment were all identical and consisted of three overlapping white discs (RGB = 255, 255, 255). The discs had a diameter of 18.75 dva diameters and were centered at the origin, 10 dva left and right of the origin, respectively. As a result, the impulse stimuli covered spatial positions of both memory items as well as the region in between. With its abrupt onset and high energy, the impulse served as a neutral, transient broadband signal that perturbed neural states by briefly activating circuits involved in encoding, thereby revealing the representational state of working memory (Stokes, 2015; Wolff et al., 2017).

The feedback was a sad/happy smiley in light gray colour and had a diameter of 5 dva. The feedback was presented at the center of the screen briefly after a response. Throughout the experiment a gray background was maintained (RGB = 128, 128, 128;, 76.3 cd/m^2^). A fixation dot formed by a black dot (0.3 dva diameter) surrounding a white circle (0.25 dva diameter), was always displayed at the center of the screen throughout a trial and participants were asked maintain their gaze fixated to this point at all times.

The experiment was programmed and run by Psychtoolbox extension for Matlab (Brainard, 1997; Kleiner et al., 2007). The stimuli were displayed on a 23.8 inch (60,452 cm on diagonal) ASUS ROG Strix XG248Q monitor with a resolution 1920 by 1080 pixels and a refresh rate of 240 Hz. The responses were collected by a regular keyboard where buttons ‘C’ and ‘M’ were pressed to indicate the direction of probe orientation. Eye position was monitored and recorded throughout a trial by a desktop-mounted Eyelink 1000 Plus at 1000 Hz sampling rate.

### Procedure

The participants completed 1600 trials divided over 10 sessions, each lasting approximately 25 minutes and conducted consecutively on the same day. In each session, 160 trials were split into 10 blocks to minimize irritation of eyes from impulse stimuli. Participants self-initiated blocks but trials within each block started automatically after each response. Participants could take breaks between blocks and sessions for self-determined durations to combat fatigue. At the start of each session participants received instructions on blinking and gaze position. Subjects were also told about the spatial position of Item 1 and Item 2, and these positions were swapped in each session. The eye-tracker was re-calibrated using a 5-dot calibration method before the beginning of each session.

During the experiment, participants sat in a dimly lit room with their heads resting on a chinrest 60 cm away from the monitor. Figure 1A illustrates a trial. Each block began when the SPACE bar was pressed, followed by a ‘Get Ready’ message presented in 12-point black Arial font for 400 ms. Next, a blank screen was displayed for 600 ms. Afterwards, the fixation dot was presented until the end of trial. After 700 ms relative to the fixation dot onset, the first memory item and a placeholder at other location each 15 dva away from the center of the screen were presented for 200 ms. An 800 ms delay followed, after which Impulse 1 was presented for 100 ms. After another 500 ms delay, Item 2 was presented for 200 ms at the other location simultaneously with a placeholder at the position of previously presented Item 1. Another 800 ms delay ensued, followed by Impulse 2 for 100 ms duration, and then a 500 ms delay. Next, a number (1 or 2) was displayed 0.25 dva above the fixation dot for 200 ms, cuing the serial position of the task-relevant item. After 800 ms delay, Impulse 3 was presented for 100 ms followed by a 500 ms delay. Finally, a probe consisting of a sinusoidal grating, was presented at the center of the screen for 200 ms. After the probe, a blank screen was presented until response. The probe was always different from the cued item. The task was to compare the orientation of the probe with the cued item. Participants responded by pressing ‘C’ or ‘M’ on the keyboard to indicate counter-clockwise and clockwise probe direction, respectively. Following their response, a smiley face (happy or sad) was displayed on the screen for 200 ms as feedback. The next trial began with the onset of the fixation cross after a random delay of 200 – 500 ms. Throughout the experiment simultaneous EEG data and eye position were monitored and recorded.

### EEG Acquisition and Preprocessing

The 64 channel EEG was recorded at 1024 Hz via BioSemi Active 2 system. The electrodes were deployed according to international 10-20 system whereas four external electrodes were placed below and above the right eye (Vertical) and lateral to the external canthi (Horizontal) to record bipolar EOG. An additional pair external electrodes was placed on mastoids to serve as a reference after recording.

The preprocessing and visual checks of the EEG data were carried out using the freely available Matlab toolboxes, Fieldtrip (Oostenveld et al., 2011) and EEGLab (Delorme & Makeig, 2004). The data was first re-referenced to the average of mastoid electrodes and it was downsampled to 500 Hz. Next, a bandpass filter (0.1 Hz highpass – 40 Hz lowpass) was applied to the EEG. ICA was used to determine components associated with eyes. After visualizing these components and their position in topographical maps, we removed blink related components for each subject individually. Next, noisy channels were identified by first checking channel-wise variance in summary statistics and then after visual inspection. These channels were replaced by spherical spline interpolation, with an average of 0.8 channels per subject. Next, data was epoched relative to stimuli onset times (−200– 1000 ms Item 1 and 2; - 200 – 600 ms Impulses 1, 2 and 3; −200 – 1000 Cue). All epochs were visually inspected for eye and muscle artefacts, drifts and noise. Trials with artefacts were removed from analyses, leading to the exclusion of 8% of trials from each epoch for each participant.

Alpha power data was obtained by band-pass filtering (8 Hz highpass – 12 Hz lowpass) continuous, non-epoched data and applying Hilbert transformation. The absolute values were epoched using the same time windows as voltage data, and the same noisy electrodes and artefact containing trials were excluded from the analyses.

### Behavioural Analyses

Behavioural analyses investigated performance in discriminating probe orientation relative to memory, using all trials. Response accuracy for all trials with the same probe deviation was averaged forming a probability distribution of correct response. Performance for probe deviations in counter-clockwise direction was then multiplied by −1, so the entire distribution reflected the probability of reporting the probe as clockwise. These calculations were completed for the cued item to reveal overall performance, as well as separately for trials where item 1, item 2 or the Left and Right item was cued and compared to the probe.

A logistic function was fitted to the observed probability of clockwise response for each subject when cued item was Item 1, 2, Item Left, or Item Right and slopes were estimated using the formula:

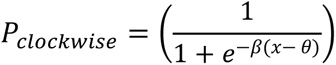

In the formula, *x* identifies probe deviation whereas the estimated *θ* is the threshold for clockwise-counterclockwise dichotomy and *β* is the slope. Fitting was performed by the *lsqcurvefit* function of MATLAB (version 2024a) where initial value for β was 1 and for θ was 0. Visual inspection indicated that logistics function captured subject level means accurately. Estimated slopes were contrasted between conditions with pairwise t-test. The confidence interval for mean slope is calculated by bootstrapping (n_perm_ = 100 000).

### EEG Analyses

#### Time-course analyses

The decoding procedure was adapted from earlier studies (Wolff et al., 2020a; 2020b). Unless otherwise stated, all multivariate analyses were conducted using data from 19 posterior electrodes (PO7, PO3, POz, Pz, P1, P3, P5, P7, P9, Oz, P2, P4, P6, P8, O1, P10, PO8, PO4, O2) based on our previous study (Kandemir & Olivers, 2024). For the time-course analyses, data in each trial were baselined by subtracting the average of −200 – 0 ms period preceding stimulus onset (e.g., memory item, cue or impulse onset). In the alpha power analyses, no baseline was used, whereas the remainder of the procedure was identical for voltage and alpha power decoding.

For decoding, first, the data was downsampled to 125 Hz to save computation time (8 ms sample spacing). Next, the data was divided into eight folds with stratified sampling to ensure that the eight orientation conditions were equivalently represented in each fold. The training set consisted of data from seven folds, used to compute a covariance matrix with a shrinkage estimator (Ledoit & Wolf, 2004). Condition specific ERP was calculated for each channel as patterns for orientation condition by averaging signal over training set trials. The number of trials in each condition was equalized by randomly sampling from the available trials to match the condition with the least trial count.

The difference between each trial in test set (remaining fold) and these patterns was calculated in Mahalanobis distance (De Maesschalck et al., 2000) with the logic that this value would be smallest if a trial belonged to one condition. The resultant distance measure was mean centered and their sign was reversed to associate higher values with higher similarity and these values are reported as similarity matrices.

To obtain a decoding accuracy value, trial specific distance measures were scaled by a cosine similarity function, reflecting radial differences between target and other orientations. The mean of this scaling operation yielded an arbitrary decoding accuracy at that time point in the trial. To mitigate variations in results due to trial selection, the entire decoding procedure was repeated 100 times, and the outputs were averaged for each participant across trials. The resulting output was smoothened using a Gaussian filter over the time dimension (kernel = 2 time points, 16 ms). Group data from all participants was then entered into statistical analyses to assess whether mean decoding was significantly different from chance level decoding obtained in null distributions (see below for how these were obtained).

To assess the generalization of the activity patterns across spatial and temporal positions, we used one of the datasets to train our algorithm and used other set as test set. In most cases contrasted conditions were not from the same trials (e.g. spatial generalization cross locations), avoiding autocorrelation. In other cases, autocorrelation posed a risk (e.g. generalization over item order across locations) in which case we used 8-fold cross-validation to sample orthogonal trials for training and test conditions.

Dynamic changes in the neural code were investigated by training on the neural data at one time point and testing it on all other time points. For this, eight-fold cross-validated decoding procedure was applied. Temporal generalization analysis was repeated ten times to account for selection bias.

#### Peak timing analyses

Cluster-based tests do not allow for a direct comparison about onset times of effects (Sassenhagen & Draschkow, 2019) Therefore we used a jackknife approach based on the peak of decoding accuracy. For this, subject means were averaged by leaving out data from one participant each time and the time-point at which this reached maximum was taken. In order to contrast peak-timing of two different decoding results, we subtracted peak decoding times and contrasted this difference against zero using a group permutation test.

#### Critical time-window analyses

When more power was required to trace memory signals within a period of time, critical time-window analyses were conducted by combining data from all 19 electrodes and time points within the time window of interest into a single vector. Time of interest was determined as 100-400 ms window relative to impulse onset based on earlier practices. This method differs from time course decoding in how the data is prepared. First, instead of using a classical baseline, the mean activity on each electrode within the critical time-window was subtracted. Then, the data within this time window was downsampled to 100 Hz by creating 10 ms long snippets using a moving average. Next, the time and channel dimensions were concatenated, yielding a single spatio-temporal pattern with 570 features. This data was decoded using 8-fold cross-validation technique, with Mahalanobis distance as the measure of similarity, just as in time-course analyses.

#### Topography

The topographical distribution of the signal was inferred from electrode groups that could significantly be decoded for memory representations using a searchlight technique (van Ede et al, 2019). First, the mean activity within the time window of interest was subtracted. Next, the data was downsampled to 100 Hz by calculating the moving average over 10 ms windows. Before decoding, data from each electrode and two of its spatially closest neighbours were pooled over time to form spatio-temporal patterns for that specific region. The 8 fold cross-validated decoding was conducted on this subset of electrodes, and the process was repeated until all electrodes were tested. This procedure was repeated 100 times to account for sampling bias. The output was then averaged over all trials and repetitions for each participant producing a single mean decoding value per electrode. Group level statistical analyses were conducted on this product.

#### Statistical analyses

Time-course and critical-window decoding output were assessed for statistical significance by comparing it to chance level decoding. For chance level decoding (null distributions), we ran the same analysis script for 1000 iterations during which condition labels (e.g. trial-specific orientation) were shuffled randomly in each time. The output of this procedure was averaged over trials. On group level, null distributions were used to estimate t-values at each time point and t-values obtained from real observations were contrasted with a cluster-based correction, using the cluster_test function (Spaak, 2015), with 0.05 as the cut-off proportion. For critical window analyses the same procedure was applied to obtain t-values for observed data and null distributions. All reported tests were two-tailed unless specifically stated otherwise. For time course data (i.e. jackknife analyses), confidence intervals for the group mean were calculated with bootstrapping (n_perm_ = 100,000).

For temporal generalization tests and searchlight analyses, we did not form null distributions this way, due to computational load, and also because these analyses were only secondary (see Supplementary Materials). Instead for temporal generalization, a null distribution was generated using the group data, by randomly flipping the sign of the mean decoding accuracy for each participant 100 000 times with a probability of 0.5. Contiguous time points exceeding a cluster-forming threshold (*p* < 0.05, uncorrected) were grouped into clusters, and the sum of their t-values was used as the cluster statistic. The maximum cluster statistic from each permutation formed the null distribution. The observed group-level cluster statistic was compared against the null distribution, and the p-value was computed as the proportion of permutations with a cluster statistic exceeding the observed value. The cut-off for statistical significance was pre-determined at 0.05. The same method was used on 2D temporal generalization plots where clusters were formed in both directions reflecting training and testing time. Output from searchlight decoding was tested with a sign-permutation test, where the sign of the data from each electrode was flipped with 0.5 probability, over 100 000 iterations to generate a null distribution. The group mean was contrasted with this distribution, the proportion it corresponded to yielding the p value for each channel while channels with *p* < 0.05 (one-tailed) were considered as electrodes significantly contributing to the memory classification.

## Results

### Behavioural Results

The overall mean response accuracy was 80.7% (sd = 6.14 %). Figure 1 show the probabilities of reporting a clockwise deviation averaged across participants, as a function of the true deviation, for serial position (Item 1, Item 2; Figure 1B) and location (Item Left, Item Right; Figure 1C), together with the best fitting psychometric function. Participants performed better when an item was shown second (Mean _slope item 1_: *β =* 0.076; Mean _slope item 2_: *β* = 0.099, *p* < 0.001). There was no difference in performance for items presented on the left versus right (Mean _slope left_: *β* = 0.88; Mean _slope right_: *β* = 0.86, *p* = 0.26). Despite the eccentricity, behavioural performance was thus quite strong. The recency effect can be attributed either to the second item interfering with the first, or the second item requiring a shorter maintenance period, and hence suffer less from decay or random noise (Broadbent & Broadbent, 1981; Schneegans & Bays, 2018; Wolff et al., 2020a).

### Decoding Results: Pre-cue phase

Here we tested for spatial generalization prior to the cue, after first establishing location-specific signals. Moreover, we assessed if such generalization occurred in the broadband signal, in alpha, or in both.

#### Spatial generalization in the broadband voltage signal

We first present the decoding of spatially specific and spatially generalized activity patterns in the broadband voltage signal. Fig. 2A (Lime colored line) shows the location-specific decoding performance averaged across left and right locations and across the first and second item. Location-specific orientation information could be successfully decoded from about 140 ms after display onset, after which decoding strength waxed for about 500 ms and then slowly waned for another 500 ms. The small inlay panel of Fig. 2A shows the decoding accuracy averaged across the significant cluster (136 - 992 ms relative to item onset, *p = 0.002*) tested against label-shuffled chance-level decoding, and which confirmed strong evidence for location-specific orientation memory [*t*(29) = 5.292, *p* < 0.001; BF_+0_ = 3690; all *one-tailed*]. In supplements-1, Figure S1A shows that the same pattern holds for each of the individual items (Item 1 and Item 2). Furthermore, similarity matrices presented in supplementary Figure S1C reflect a parametric-looking relationship as the similarity was reduced as a function of angular differences between memoranda, which provides further evidence that decoded signal is indeed associated with orientation memories. For completion we also provide temporal generalization analyses plus the scalp origin of these signals (Figures S1G and S1H). The relatively slow development of the orientation signal is consistent with what has been observed before for peripherally presented items and differs from typical patterns reflecting centrally presented items (Kandemir & Olivers, 2024). We will return to this in the General Discussion.

**Figure 2.**
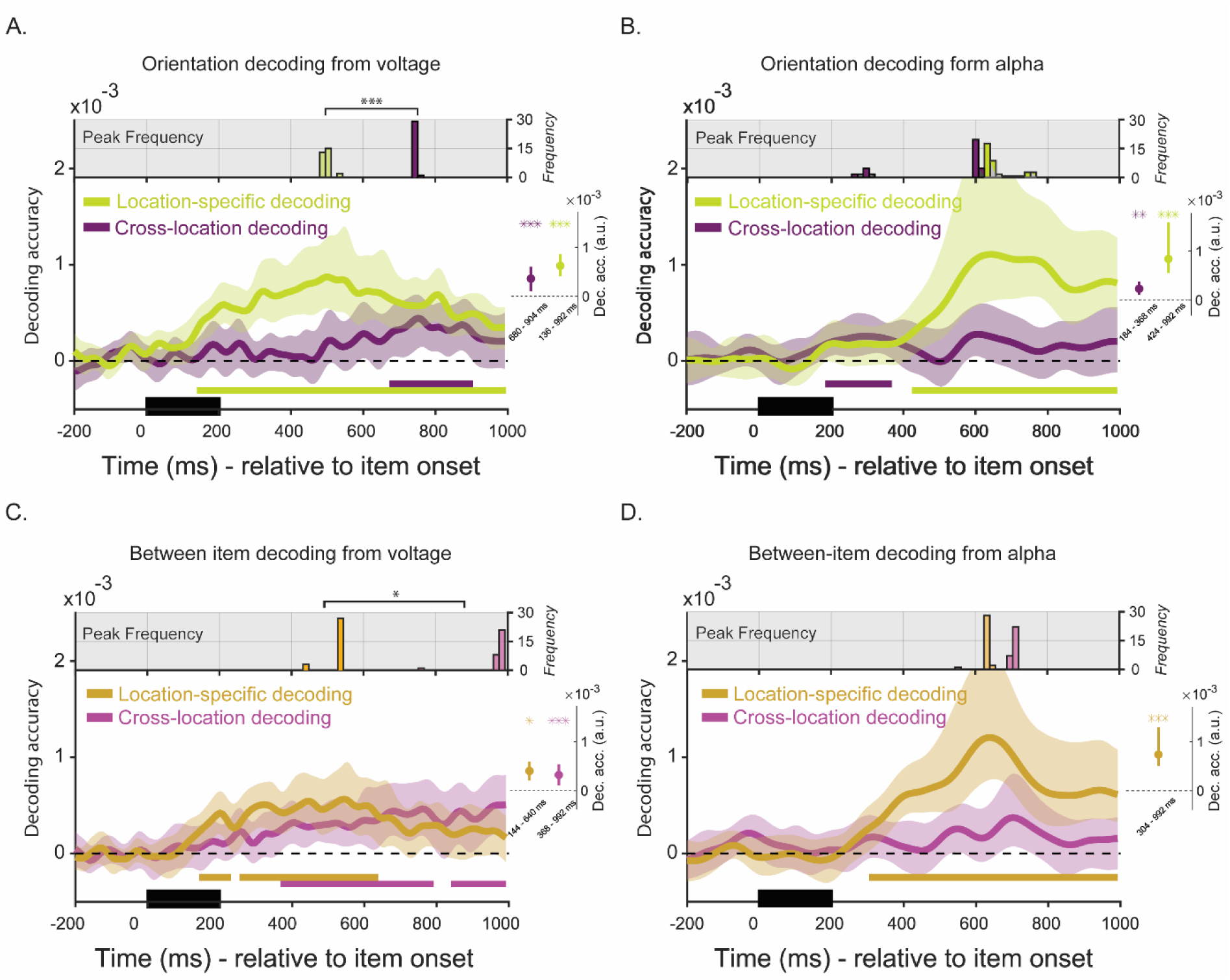
Time course of location-specific and cross-location decoding of orientations following the presentation of the memory items, from both voltage (A) and alpha power (B) data., Panels (C) and (D) also show location-specific and cross-location decoding for voltage and alpha, but now the classifier was trained and tested across temporal positions. *Note.* Orientation decoding and the generalization of neural patterns across spatial and temporal positions as a function of time. **A-B.** Time-course location-specific (lime) and cross-location (plum) decoding of Item 1 and 2 (averaged) using voltage and alpha power data. **C-D.** Time-course location-specific (yellow) and cross-location (pink) generalization of orientations between spatial and temporal positions using voltage and alpha power data. **A-B-C-D.** Solid colored lines mark mean decoding performance over time. Colored shadings mark 95% C.I. Colored bars below designate time window with statistically significant decoding (p < 0.05) relative to chance level decoding. Black bars starting at time 0 marks presentation duration of memory items. Boxplots on the right show mean decoding accuracy within statistically significant period and 95 % C.I. of the mean. Asterisks reflect statistical significance (*** = p < 0.001, ** = p < 0.01, * = p < 0.05). Top panels show frequency distribution of time points with peak decoding, according to jackknife procedure. Asterisks show time difference with statistical significance.

To assess spatial generalization, we computed the average decoding performance *across* locations, by training on one location and testing on the other, again averaged for the first and second item (Figure 2A; Plum colored line). The spatially generalized signal revealed a different time course compared to the location-specific signal. Notably, spatial generalization emerged in the later part of the pre-cue phase, (680 – 904 ms cluster relative to item onset, *p* = 0.002), which was also highly reliable when averaged across the cluster and tested against label-shuffled baseline [*t*(29) = 4.794, *p* < 0.001; BF_+0_ = 1070]. Here too, similarity matrices (Supplements −1, Fig S1D) showed a parametric looking relationship between neural codes for memoranda further indicating that decoding was associated with orientation information. While the different timings of the clusters are suggestive, cluster-based permutation analyses are not suitable for assessing timing differences (Sassenhagen & Draschkow, 2019), and so we adopted a jackknife procedure to statistically assess temporal differences between conditions in peak decoding accuracy (see Figure 2A, top panel for the frequency distribution of peak decoding time points). This revealed a clear temporal difference between time points at which the location-specific and cross-location neural codes peaked (Location-specific M = 503 ± 12.0 ms; Cross-location M = 752 ± 1.5 ms; *p* < 0.001).

We then assessed if this pattern could be confirmed when decoding across temporal positions, by training the classifier on Item 1 and testing it on Item 2, and vice versa. We again did this within the same location (location-specific decoding) and cross-locations (testing for spatial generalization). Figure 2C reveals a pattern similar to that of Figure 2A, with cross-location decoding emerging later than location specific signals. (Fig. 2B, location specific, yellow, 144 - 232 and 256 - 640 ms relative to item onset, *p* = 0.006 and *p* = 0.002; cross location, pink, 368 – 608 ms and 640-792 ms relative to item onset, *p* = 0.004; and 840 - 992 ms relative to item onset, *p* = 0.006). In both cases orientation information also surpassed label-shuffled baselines [see the small inlay panel in Fig. 2C, location specific, yellow, *t*(29) = 4.523, *p* < 0.001; BF_+0_ = 541; pink, cross location, *t*(29) = 3.299, *p* = 0.001; BF_+0_ = 28.88; all *one-tailed*]. Supplementary Fig S1E provides the parametrically changing similarity matrices. To assess timing differences we again performed a jackknife analysis, and this again revealed that location-specific information peaked earlier than spatially generalized information (Figure 2C, top panel, location specific M = 533 ± 31.6 ms; cross location, M = 977 ± 42.6 ms, *p* = 0.03).

Collectively then, the voltage decoding results indicate a transformation from spatially specific representation into a spatially generalized representation. Importantly, this spatial generalization occurs before the cue, and thus before the participant knows which item will become relevant for report.

#### Little to no spatial generalization in the alpha signal

We performed the same analyses on the alpha frequency signal. Here the dynamics were quite different. Figure 2B shows how location specific orientation information appeared relatively late (Lime color, 424 – 992 ms relative to item onset, *p* = 0.002; t(29) = 3.668, *p* < 0.001; BF_+0_ = 67.64; *one-tailed*), while cross location decoding hardly materialized bar for relatively weak early cluster (Plum color, 184 – 368 ms relative to item onset, *p* = 0.014; *t*(29) = 3.09, *p* = 0.002; BF_+0_ = 18.06). Figure S1B in Supplements-1 shows decoding performance for the individual items (Item 1 & Item 2), and the parametrically changing similarity matrices. A jackknife analysis on peak decoding time points for generalized and location specific signals did not reveal a reliable timing difference (Fig. 2B, top panel; location-specific: M = 657 ± 40 ms; cross-location: M = 533 ± 137.8 ms, *p* = 0.87), but we again emphasize that for this analysis there was little generalized signal to begin with.

This pattern was confirmed when decoding across temporal positions (i.e. between item 1 and 2). Figure 2D revealed reliable location specific orientation decoding (location specific, yellow, 304 - 992 ms relative to item onset, *p* = 0.002; *t*(29) = 4.109, *p* < 0.001; BF_+0_ = 194.9, *one-tailed*) but not across spatial locations. Jackknife procedure could not reveal a significant difference in peak decoding times (Figure 2D, top panel, location specific: M = 638.4 ± 4.4 ms; cross location, M = 697.8 ± 28.0 ms, *p* = 0.70), while again pointing out the caveat that it is difficult to estimate the peak of a largely absent generalized signal.

Thus, the results reveal a clear dissociation of relatively early location-specific orientation information in the voltage signal, which then generalizes during the second half of the delay period, from about 400-600 ms onwards. This later time window is also when the location-specific alpha signal emerged, which however showed no sign of generalization.

#### Control for eye movement confounds

To check for any confounding signals form eye movements in the reported analyses, we repeated the above decoding analyses using gaze position coordinates as recorded by the eye tracker (see Figs S2A-C in Supplementary Material 2). We observed no reliable location specific decoding, nor any spatial generalization in the eye signal, and thus conclude that the observed EEG decoding patterns are not driven by eye movements.

#### Impulse-related activity

As activity tends to wane during VWM delay periods, we also included an impulse stimulus during the delays prior to selection, as such impulses have been shown to revive mnemonic patterns. Figure 3A shows reliable location-specific activity patterns post impulse [location-specific, lime, 0 – 592 ms relative to impulse onset, *p* = 0.002; *t*(29) = 4.25, *p* < 0.001; BF_+0_ = 273, *one-tailed*, when cluster-wide average is tested against shuffled labels baseline), as well as spatially generalized activity (cross-location, plum, 176 – 472 ms relative to impulse onset, *p* = 0.002; *t*(29) = 2.98, *p* = 0.003, *one-tailed*; BF_+0_ = 14.39). The supplementary material shows decoding performance for the individual items, as well as the parametric tuning matrices and topographical distribution (Supplements −3, Figs S3A to D).

**Figure 3.**
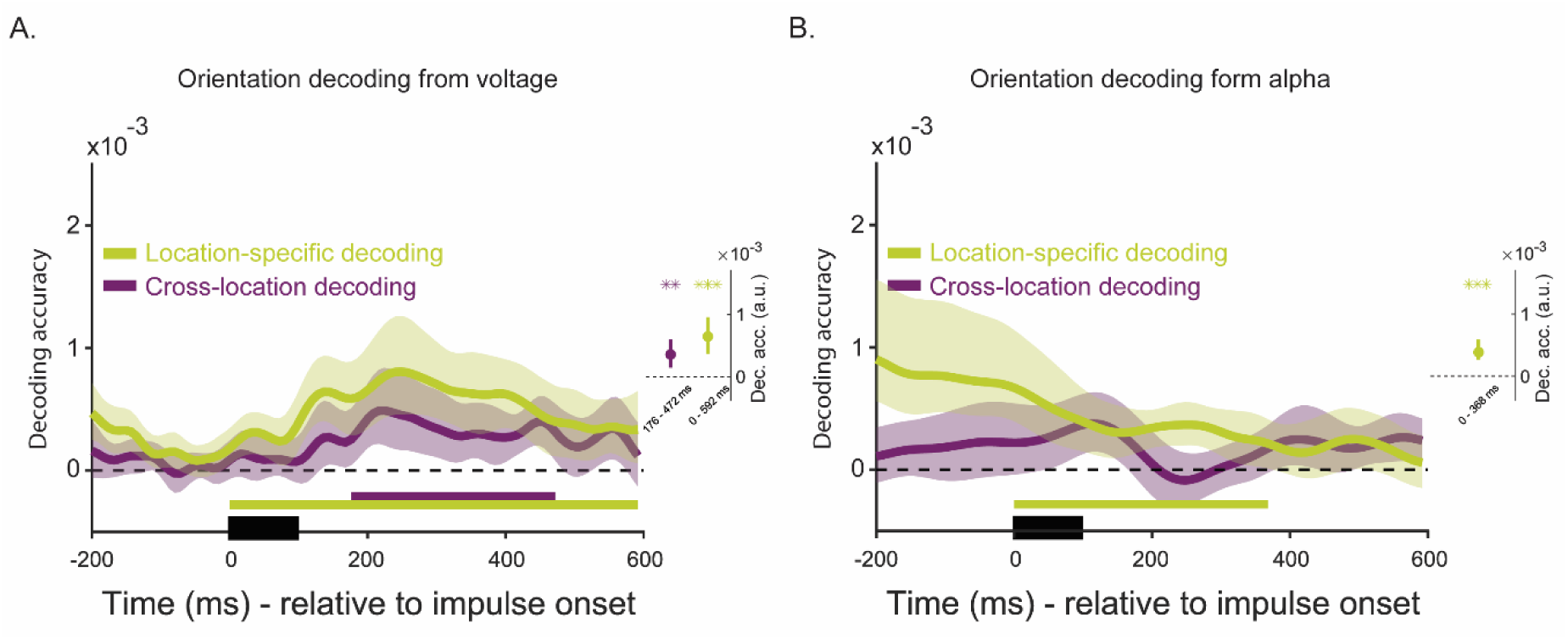
Time course of location-specific and cross-location decoding of orientations from voltage (A), and alpha power data following the presentation of the impulse during the first delays, prior to the cue. *Note.* Orientation decoding after the presentation of impulse 1 and 2. **A.** Time-course location-specific (lime) and cross-location (Plum) decoding for orientations, averaged over Item 1 and 2 as well as Impulse 1 and 2 from voltage data. **B.** Time-course location-specific (lime) and cross-location (Plum) decoding for orientations, averaged over Item 1 and 2 as well as Impulse 1 and 2 from alpha power data **A-B.** Solid colored lines represent mean decoding performance as a function of time and the surrounding shades mark 95 % C.I. Colored bars mark time window with above chance level decoding with statistical significance (*p* < 0.05). Black rectangle marks the duration of impulse presentation. Boxplots on the right show mean decoding accuracy over time window with statistical significance and 95 % C.I. of the mean. Asterisks reflect statistical significance (*** = *p* < 0.001, ** = *p* < 0.01, * = *p* < 0.05).

Figure 3B shows the same post-impulse analysis but here for the alpha-band signal. Location-specific orientation decoding was statistically significant before impulse and we observed a significant cluster when we focused on the period following impulse onset (Lime color, 0 – 368 ms relative to impulse onset, *p* = 0.002; t(29) = 5.33, *p* < 0.001; BF_+0_ = 4220; *one-tailed*). Contrary to the voltages, the presentation of the impulse led to a reduction in decoding accuracy (M dec. acc. −200 – 0 ms vs. M dec. acc. 100 – 300 ms relative to impulse onset, *t*(29) = 2.53, *p* = 0.009, *one-tailed*), eventually obliterating it, consistent with what we observed in our previous study (Kandemir & Olivers, 2024). Note that this does therefore not mean that alpha could not in principle contain orientation information at this stage during the delay, but that it was likely obliviated by impulse-evoked signals. On the other hand, cross-location decoding (Fig. 3B, Plum color) was not significant in alpha band prior to impulse nor did the impulse presentation increase it.

### Post-cue phase

Figure 4A shows the location-specific orientation decoding of the broadband voltage signal for cued and uncued items. As can be seen, and to our surprise, we could not decode orientation information from these data, even for the cued item. Figure 4C shows the decoding across locations, and it is clear that we found no spatially generalized signal either, whether for cued or uncued items.

**Figure 4.**
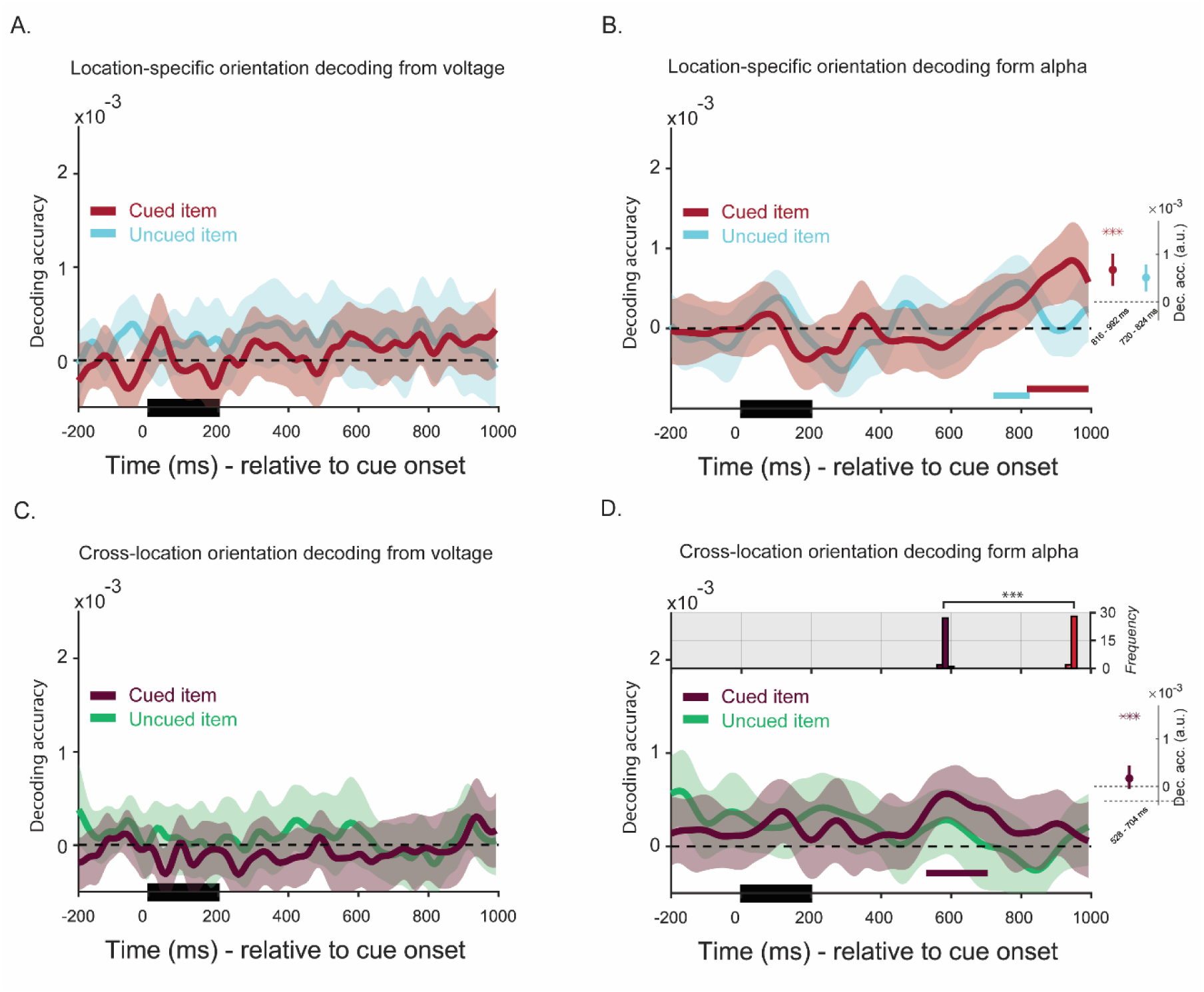
Time course location-specific and cross-location decoding of cued and uncued item relative to cue from voltage (A and C) and alpha power data (B and D),. *Note.* Decoding task-relevant cued and task irrelevant, uncued item after cue presentation. **A.** Time-course location specific decoding for cued (red) and uncued (cyan) item from voltages, averaged over spatial and temporal positions relative to cue onset. **B.** Time-course location specific decoding for cued (red) and uncued (cyan) item, from alpha power, averaged over spatial and temporal positions relative to cue onset. **C.** Time-course cross location decoding for cued (maroon) and uncued (green) item from voltages, averaged over temporal positions relative to cue onset. **D.** Time-course cross location decoding for cued (maroon) and uncued (green) item, from alpha power, averaged over temporal positions relative to cue onset. **A-B-C-D.** Solid colored lines represent mean decoding accuracy whereas shadings mark 95 % C.I. of the mean. Solid colored bars mark the period with statistically significant decoding relative to chance level, according to cluster corrected test (*p* < 0.05). Black bars starting at time 0 mark the stimulus (here cue or impulse 3). Boxplots on the right show mean decoding accuracy for each condition, averaged over the window with statistically significant decoding and 95 % C.I. of the mean. **F-G.** Large dots mark mean values whereas colored bars show 95 % C.I. The lower and upper borders of the boxes indicate 25th and 75th percentiles whereas whiskers mark +/− 1.5 IQR. **A-B.** Asterisks reflect statistical significance (*** = *p* < 0.001, ** = *p* < 0.01, * = *p* < 0.05).

Figure 4B shows location-specific orientation decoding of the alpha signal. Here we could observe statistically significant orientation information for the cued item at the end of the epoch (red, 816 - 992 ms relative to cue onset, *p* = 0.002; *t*(29) = 3.97, *p* < 0.001; BF_+0_ = 140), and to a lesser extent but still momentarily reliable for the uncued item (cyan, 720 - 824 ms relative to cue onset, *p* = 0.026), although this was not reliable when tested against a label-shuffled baseline (*t*(29) = −0.579, *p* = 0.72; BF_+0_ = 0.13)). Interestingly, a spatially generalized signal for the cued item emerged already emerged before that in the alpha band signal (Fig. 4D, cued item, maroon, 528- 704 ms relative to cue onset, *p* = 0.006; *t*(29) = 3.79, *p* < 0.001; BF_+0_ = 9176; not for the uncued item, green, *n.s.*). A jackknife analysis indeed confirmed that generalized and location specific signal peaked at distinct time points (cross-location M = 587.2 ± 5.4 ms; location-specific, M = 944.3 ± 3.3 ms, *p* < 0.001). It may be that in the reverse cascade of retrieving the original information the re-activation of the voltage-related signal is dependent on the alpha signal, and that the post-cue period was simply not long enough to observe re-activation of the voltage signal. While speculative, the next analysis provides some support, as the voltage signal did return after the impulse.

During the post-cue delay, another impulse signal was presented. The orientation of the cued item identity could be successfully decoded from the voltage data just after this impulse (Figure 5A, Cued item, red, 200 - 272 ms and 328 – 384 ms relative to Impulse 3 onset, *p* = 0.016; *t*(29) = 2.93, *p* = 0.003; BF_+0_ = 12.80), while there was no reliable trace of the uncued item. As at this stage signals were generally weaker, we next pooled over electrode and temporal space, by combining data from all 19 electrodes within 100-400 ms time-window into a single vector, and decoded cued and uncued item separately per location (Figure 5B). This confirmed significant above-chance location-specific orientation decoding for cued items (Cued item, red, 100 - 400 ms relative to Impulse 3 onset, *p* = 0.01), but not uncued (cyan, ns). Time course cross-location decoding was statistically significant for the cued item although this effect was very brief and weak (Figure 5C, Cued item, maroon, 216 - 256 ms relative to Impulse 3 onset, *p* = 0.036; *t*(29) = 2.45, *p* = 0.01; BF_+0_ = 4.89), but not for the uncued item (Uncued item, green, *n.s.*). Pooling over electrode space within 100-400 ms period, we observed spatial generalization for the cued item as well (Cued item, maroon, 100-400 ms relative to Impulse 3, *p* = 0.018), which was not the case for the uncued (green, ns) while the difference between cued item and uncued items was statistically significant (*p* = 0.016, *one-tailed*). A comparison of the location-specific and spatially generalized decoding of the cued item did not reach statistical significance. This then raised the question whether the location-specific and spatially generalized codes coexisted or observers adopted one format on one trial and the other format on another trial. To test this we correlated decoding accuracy across trials between the general and the localized codes. We observed a weak positive correlation, which was statistically significant against zero (Mean Fisher transformed *r* = 0.016, sd = 0.037, *p* = 0.029). Such a correlation was not present for the uncued item (Mean Fisher transformed *r* = 0. 004, *p* = 0.60), which served as control in this analysis. The positive correlation is more consistent with coexistence than with a trade-off between location specific and location general codes.

**Figure 5.**
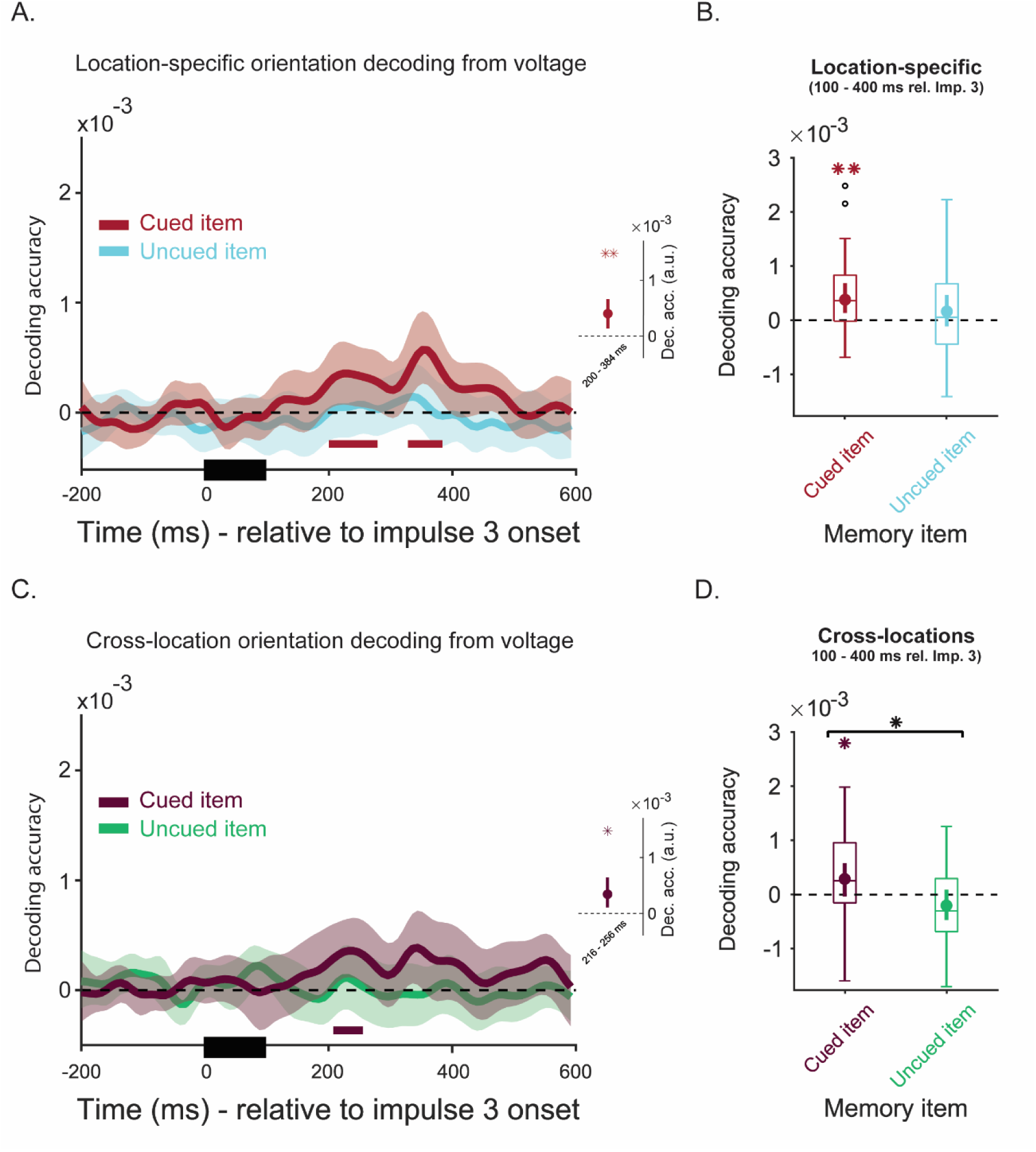
Time course location-specific and cross-location decoding of cued and uncued item relative to impulse 3 from voltages (A and C), alongside trial wise location specific (B) and cross location decoding (D) from voltages when data was pooled over electrode and temporal space (100-400 ms time-window). *Note.* Decoding task-relevant cued and task irrelevant, uncued item after impulse 3. **A.** Time-course location-specific decoding for cued (red) and uncued item (cyan) relative to impulse 3 onset. **B.** Boxplots displaying location-specific decoding accuracy for the cued (red) and uncued item (cyan) with data combined over electrode and temporal space within 100 – 400 ms window relative to impulse 3. **C.** Time-course cross-location decoding for cued (maroon) and uncued (green) item relative to cue onset. **D.** Boxplots displaying cross-location decoding accuracy for the cued (maroon) and uncued item (green) with data combined over electrode and temporal space within 100 – 400 ms window relative to impulse 3. **A-C.** Solid colored lines represent mean decoding accuracy whereas shadings mark 95 % C.I. of the mean. Solid colored bars mark the period with statistically significant decoding relative to chance level, according to cluster corrected test (*p* < 0.05). Black bars starting at time 0 mark the stimulus (here cue or impulse 3). Boxplots on the right show mean decoding accuracy for each condition, averaged over the window with statistically significant decoding and 95 % C.I. of the mean. **B-D.** Large dots mark mean values whereas colored bars show 95 % C.I. The lower and upper borders of the boxes indicate 25th and 75th percentiles whereas whiskers mark +/− 1.5 IQR. Asterisks reflect statistical significance (*** = *p* < 0.001, ** = *p* < 0.01, * = *p* < 0.05).

## Discussion

Information in visual working memory spatially extends beyond the location of the encoded stimulus, suggesting a transition from location-specific representations to a common code. Previous work has demonstrated spatial generalization both using fMRI- and EEG-based methods (Bae & Luck, 2018; Ester et al., 2009; Fukuda et al., 2016; Kandemir & Olivers, 2024; Kandemir et al., 2024a). However, it has remained unclear what the dynamics are of this generalization and what process or processes it is associated with. In previous studies only a single memorandum was presented, or generalization was only measured after participants had selected a single time from memory, which leaves the question whether generalization is related to the original sensory features (here orientation), or a shared mapping sensory input onto a common spatial abstraction or output-related representation spatially, or both (Bae & Luck, 2018, 2019; Bae & Chen, 2024; Cochrane & Green, 2023; Duan & Curtis, 2024; van Ede et al., 2019; Fahrenfort et al, 2017; Henderson et al., 2022; Kandemir et al., 2024a; Kwak & Curtis, 2022; Mostert et al., 2018; Panichello & Buschman, 2021). In the current study we sought to test which processing stage comes with generalization by instructing observers to remember two patterns on each trial and only indicate later (using a retro-cue) which of the two would be relevant for report.

We observed a clear transition from location-specific activity patterns to spatial generalization already during the maintenance period prior to the selection cue. Moreover, this spatial generalization was independent of presentation order during encoding, as it also occurred between first and second items in the sequence. The location-specific information peaked around 400 ms, which was about the time at which the spatially generalized information only started to emerge, in turn reaching maximum levels around 800 ms post display onset. Both these location-specific and generalized voltage signals then re-emerged after the first impulse (but still prior to the cue). It is noteworthy that that at 140 ms, even the location-specific orientation information emerged relatively late when compared to typical EEG studies of visual working memory, which tend to show early and strong onset-related transients in decoding that are typical for stimulus-related activity (e.g. King & Wyart, 2021; Kandemir & Akyürek, 2023; Kandemir et al., 2024b; Myers et al., 2015; Wolff et al., 2017). The crucial difference is that we presented stimuli rather peripherally, at 15° eccentricity, while previous EEG studies of visual working memory have used more centrally presented stimuli. We observed a similarly slow emerging pattern in our previous study where we presented stimuli at the same eccentricity (Kandemir & Olivers, 2024). In contrast, for foveally presented stimuli we did observe strong initial information. We hypothesize that the inability to decode the early sensory signals arises from the folding of primary visual cortex, with the periphery being represented inside the calcarine fissure (Seki et al., 1996), resulting in less detectable dipoles. The important implication is that the slower signal observed here must reflect a different signal than is initially triggered by the stimulus, and we argue that it likely represents feedback activity higher up the visual hierarchy, sustaining the actual memory.

What is the nature of the spatially generalized representation? While we have no definitive evidence, we propose that it is largely sensory in nature, for several reasons. The first reason is that by nature of the trial design, during the first maintenance period it was not yet known which item would become relevant and serve report. Furthermore, the actual response could not be meaningfully prepared until the end of the trial, as it depended on the probe orientation. Thus, it is unlikely that the generalized pattern reflected response preparation. The second reason is that the generalization occurred in the broadband voltage signal (i.e. evoked potentials). Previous work has provided evidence that information extracted from this signal is more consistent with the actual feature information being presented (typically orientation, as was also the case here; Bae & Luck, 2018; 2019; Fahrenfort et al., 2017; Kandemir & Olivers, 2024). This becomes particularly relevant in combination with the third reason, namely that prior to the cue, the generalization did *not* occur within the alpha signal, a signal that has been associated with spatial derivatives of the stimulus that observers then attend to (Bae and Luck, 2018; Bae & Luck, 2019; Bae & Chen, 2024). We note though that the fact that the alpha-band signal did not show spatial generalization does not mean that alpha, and the process it reflects, plays no role in such generalization. In fact, it is quite striking that the spatially generalized voltage signal emerges around the time of the emerging location-specific alpha signal. Although we have no conclusive evidence, this may mean that the two signals are related. For example, the generalization of the representation (as reflected in the voltage) could be the consequence of spatially attending to the memorandum (as reflected in the alpha).

A generalized sensory representation is further supported by evidence from neuroimaging work by others (Ester et al., 2009; Jehee et al., 2011; Serences & Boynton, 2007), which has shown that spatially generalized patterns can be obtained from early visual cortex. We emphasize once more though that while converging evidence points towards pre-report sensory representations, we cannot directly determine their nature and thus “how sensory” they are. Moreover, even if we assume that these generalized representations indeed reflect sensory traces, several options remain. One option is that feature representations spread from their localized representation to a global pattern of feature activity across the visual cortex (Ester et al., 2009; Fukuda et al., 2016; Serences & Boynton, 2007). An alternative is that the representations of the peripheral memoranda become centralized, transferred to areas normally representing foveal vision (Chambers et al., 2013; Weldon et al., 2016; Williams et al., 2008). This too would result in a common code, and we cannot determine from our data which of these options – globalization of fovealization – would hold true (if at all).

Our current findings contrast with our previous study (Kandemir & Olivers, 2024), where we did observe clear generalization in alpha-band oscillations during the delay after item presentation, but not in the voltage signal. We believe this dissociation is meaningful: In our previous study there was only one stimulus to remember, and this stimulus also directly dictated the prospective response in a one-to-one fashion, as the memory test required observers to push a joystick in the direction of the orientation. Hence the memorandum could in principle be directly re-coded into a common spatial or even response-related signal, which was then reflected in alpha. These signals may then have obscured or obfuscated any generalized voltage-driven activity patterns. In the current study no such common spatial or response code needed to be prepared prior to the cue.

After the cue, there was no longer an active orientation trace in the broadband voltage signal, although it could be evoked again by the impulse stimulus (both spatially specific and generalized). There was still evidence for alpha-based information, this time including generalized alpha information. This is generally consistent with the framework that after selection of a task-relevant item other types of representation than the pure sensory one become more important, presumably attention- or response-related ones. Unexpectedly though, the spatially generalized pattern emerged before the spatially specific information, which may reflect a process where observers first attend to a generalized representation as created prior to the cue, but then eventually return to the original, localized memory, perhaps because the original trace contains more or more precise information. However, we hasten to point out that signals were generally weak in this stage and so any conclusions remain very speculative. Further research will be needed.

We conclude that spatial generalization within VWM emerges within 400 to 800 ms, starting during what are likely sensory, pre-report stages of processing. After determining which items is task-relevant, additional generalization may emerge from shared attention and response processes, depending on whether the task allows for a common mapping (spatial or otherwise).

## Data Availability

Processed and anonymized EEG and behavioural data reported in this manuscript, as well as MATLAB scripts and their output can be accessed via this link: (https://osf.io/jcg5k/?view_only=092daf47d6ea499785f08c1d382157af)

## Acknowledgements

This project was funded by the Dutch Organization for Scientific Research (NWO; grant 453-16-002, to C.N.L.O.). We thank Ottavia Frangini for her help as a lab assistant during data collection and pre-processing.

