## Supplementary document 1, 2 and 3 for "Spatial generalization of peripherally encoded memories emerges before selection for report"

**Supplements -1**

Individual orientations could be successfully decoded shortly after display onset (Figure S1A, Item 1, dark blue, 144 ms – 952 ms, $p = 0.002$; Item 2, orange, 136 ms – 992 ms, $p = 0.002$) for which the temporal mean was also significantly above chance level (Fig. S1A, right panel, dark blue, Item 1, $t(29) = 4.25$, $p < 0.001$; $BF_{+0} = 274$; orange, Item 2, $t(29) = 4.56$, $p < 0.001$; $BF_{+0} = 598$; all *one-tailed*). Similarity matrices are formed by sign reversed, mean centered Mahalanobis distances ordered relative to trial specific item in each trial, averaged over trials and subjects. Brightness indicates higher similarity whereas darker colors mark dissimilarity. Gradual change in similarity measure is considered evidence that neural signal varies as a function of angular differences. Below we presented representational similarity matrices for location specific decoding of Item 1 and 2 from voltage data (Fig. S1A),

When alpha power was decoded for Item 1 and 2, memory signal slowly emerged 400 ms after display onset with increasing strength (Figure S1B, Item 1, dark blue, 512 ms – 992 ms, $p = 0.002$; Item 2, orange, 408 ms – 840 ms, $p = 0.002$ and 888 - 992 ms, $p = 0.032$). Temporal averages of decoding accuracy for item 1 and 2 within the significant clusters were significantly higher than chance-level (Fig. S1B, right panel, dark blue, Item 1, $t(29) = 3.93$, $p < 0.001$; $BF_{+0} = 119{,}5$; orange, Item 2, $t(29) = 3.38$, $p < 0.001$; $BF_{+0} = 34.65$; all *one-tailed*), reflecting a robust location specific trace in alpha band. Parametric-looking relationship observed on similarity matrices provided further evidence that alpha band decoding was associated with orientation memories.

Other similarity matrices are also presented, namely cross location and location specific decoding from voltages (Fig. S1C) and alpha power (Fig. S1D.), cross location and location specific decoding across temporal positions from voltage (Fig. S1E) and alpha power (Fig. S1F), respectively.

Temporal dynamics were investigated by training data at one point and then testing it on all available time points in order to assess changes in neural code for orientations. The output was averaged over item 1 and 2 and tested for statistical significance by using non-parametric permutation test with cluster correction. Null distribution was formed by flipping the sign of data with 0.5 probability over 10000 iterations and observed data was contrasted with this. Cluster correction ensured multiple comparisons did not inflate false positives.

Temporal generalization matrix revealed that while the initial approximate 400 ms period came with significant above-chance decoding at the diagonal (Figure S1G, where gray outlined regions indicate significant decoding above zero, $p < 0.05$), there was little in terms of off-diagonal signals, indicating that this early phase was characterized by a dynamic activation pattern. During this early phase, the topographical distribution of the decoded memory signal was strongly lateralized according to stimulus location, as was confirmed by searchlight analyses (Figure S1H, 100-400 ms window, all marked electrodes with decoding above zero, $p < 0.05$, one-tailed), consistent with a location-specific signal. In the later part of the epoch the signal became more stable, as it now generalized across time. During this phase, the topographical distribution was also more centralized on the posterior scalp (Figure S1H, 500-800 ms window, all marked electrodes with decoding above zero, $p < 0.05$, one-tailed), consistent with the possible emergence of a spatially generalized code – which we will address next.

**Figure S1**

*Time course location specific decoding for Item 1 and 2 averaged over locations with similarity matrices (SMs) and cross locations using voltage (A) and alpha power (B); similarity matrices for location specific orientation decoding averaged over items using voltage (C) and alpha power (D); SMs showing generalization between Items 1 and 2 across serial positions using voltage (E) and alpha power (F); temporal generalization of memory representations averaged across Items 1 and 2 (G); and topographical distributions of memory-related signals derived from voltage data during early (100–400 ms) and late (500–800 ms) time windows (H). All analyses are time-locked to the onset of Items 1 and 2.*

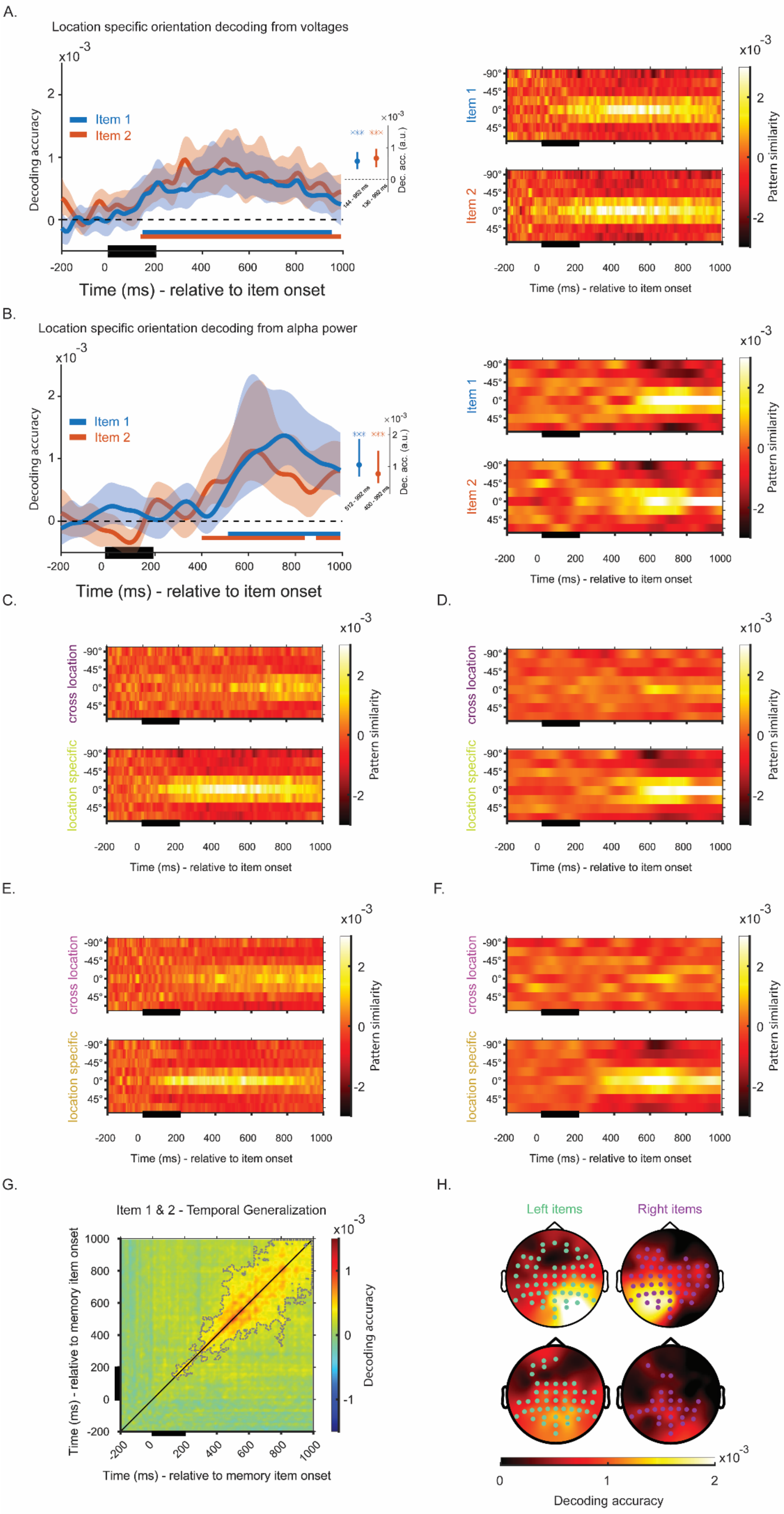

A.
Location specific orientation decoding from voltages
Decoding accuracy
Item 1
Item 2
Time (ms) - relative to item onset
Dec. acc. (a.u.)
144 - 992 ms
136 - 992 ms
Pattern similarity
B.
Location specific orientation decoding from alpha power
512 - 992 ms
400 - 992 ms
C.
D.
cross location
location specific
E.
F.
G.
Item 1 & 2 - Temporal Generalization
Time (ms) - relative to memory item onset
Decoding accuracy
H.
Left items
Right items
Decoding accuracy

*Note.* **A-B.** Time-course decoding for Item 1 (dark blue), and 2 (orange) when location differences are concerned, using voltage and alpha band data and representational similarity matrices forItem 1 and 2. Solid colored lines mark mean decoding over time. Colored shadings mark 95% C.I. Colored bars designate time window with statistically significant decoding ($p < 0.05$) relative to chance level decoding. Boxplots on the right show mean decoding accuracy averaged over time window with statistical significance and 95 % C.I. of the mean. Asterisks reflect statistical significance (*** = $p < 0.001$, ** = $p < 0.01$, * = $p < 0.05$). **C.** Representational similarity matrices for cross location and location specific decoding of voltage data for memory items averaged across item 1 and 2. **D.** Representational similarity matrices for cross location and location specific decoding of alpha power data for memory items averaged across item 1 and 2. **E.** Representational similarity matrices for generalization of item 1 and 2 across temporal positions with cross location and location specific decoding on voltage data. **F.** Representational similarity matrices for generalization of item 1 and 2 across temporal positions with cross location and location specific decoding on voltage data. **A-B-C-D-E-F.** Black bars along the time axis mark the period with item presentation. Brightness indicates higher similarity and gradual change in similarity with angular difference indicates memory representations change as a function of orientation space. **G.** Temporal generalization for memory items, averaged across serial positions. Initial period from 100 to 400 ms reveal a different temporal pattern in comparison to later period. Areas bordered by dotted lines show the time-zones with above zero decoding with statistical significance ($p < 0.05$). **H.** Topographic distribution of decoding, revealing electrode contributions for memory items on Left and Right during the initial 100-400 ms window as well as late 500-800 ms window, averaged across serial position, according to searchlight analysis. Brightness reflects the strength of decoding. Colored dots mark electrodes with above zero decoding accuracy with statistical significance ($p < 0.05$, one-tailed).

### Supplements -2

We decoded item identity using gaze position recorded by the eye-tracker. First, the data was downsampled to 500 Hz and epoched relative to item 1 and 2 onset (- 200 ms to 1000 ms). The missing data due to blinks was filled in by first calculating mean x and y coordinates before and after a blink, using preceding or subsequent time points (1, 5 or 10 timepoints when available). The missing samples were then filled with linearly and equally spaced values generated between the mean values. If eye data was completely missing, that trial was excluded. The same data was also decoded with 1000 iterations during which condition labels were shuffled randomly, in order to generate a null distribution with chance level decoding. Observed data was contrasted with null distribution using cluster correction.

When gaze position was decoded for item 1 and 2 separately for each location (Fig. S2A, blue, Item 1; orange, Item 2), cross locations (S2B, black, Item 1&2 – location specific, gray, Item 1 & 2, cross locations) and cross temporal positions, we observed no difference from chance level decoding in either condition. We concluded that our EEG decoding was not driven by eye movements.

**Figure S2**

*Time course location specific decoding for Item 1 and 2 averaged over locations (A) cross locations (B) and temporal positions (C) from gaze position.*

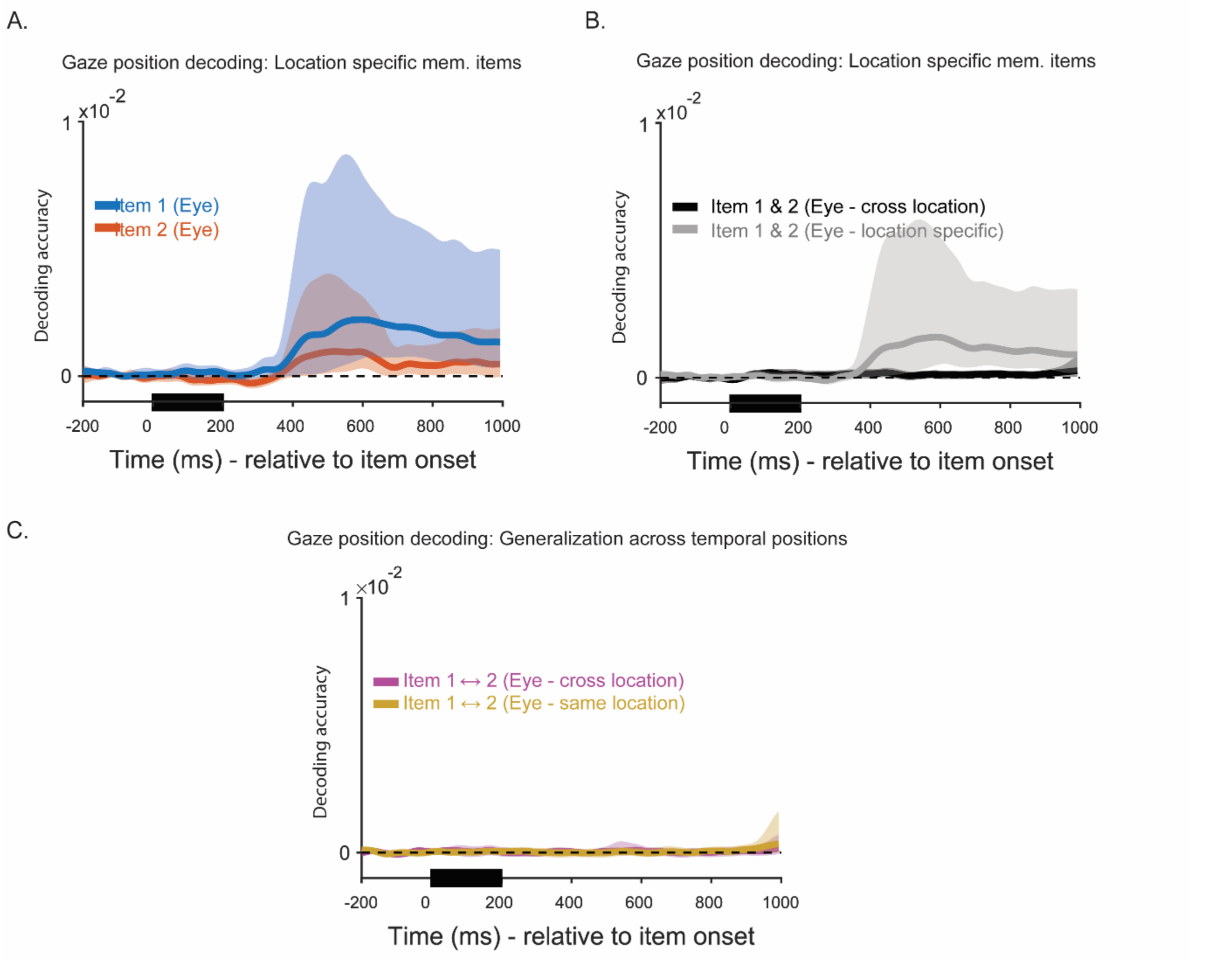


*Note.* **A.** Time course mean decoding of item 1 (dark blue) and 2 (orange) from gaze position (x, y coordinates measured by eye tracker) specific for each item location, averaged over temporal order . **B.** Time course mean decoding for spatial generalization of memory items (black) and location-specific decoding (gray), averaged over item 1 and 2 from gaze position (x, y coordinates measured by eye tracker). **C.** Time course mean decoding for generalization over temporal positions location specific (yellow) and cross location (pink) from gaze position (x, y coordinates measured by eye tracker). **A-B-C.** Colored solid lines represent the mean across all participants and trials. The shaded areas mark 95 % C.I. Solid black bars below mark the time-window during which memory items were presented. No significant clusters were detected when comparing decoding accuracy to chance level decoding obtained by shuffling labels, over 1000 iterations.

### Supplements -3

During maintenance we observed successful decoding of both memoranda individually, after the presentation of impulse 1 and 2 (Figure S3A, Item 1, dark blue, 0- 64 ms, $p < 0.008$ and 88 ms – 456 ms relative to Impulse 1 onset, $p = 0.002$; Item 2, orange, 104 ms – 592 ms relative to Item 2 onset, $p = 0.002$). Averaging over the time window with statistical significance, both items were reliably higher than chance level decoding (Fig. S3A, right panel, dark blue, Item 1, $t(29) = 3.87$, $p < 0.001$; $BF_{+0} = 108$; orange, Item 2 , $t(29) = 3.09$, $p = 0.002$; $BF_{+0} = 17.96$; all *one-tailed*). Similarity matrices for location specific decoding of Item 1 and 2 revealed a parametric looking relationship between orientations. This was also the case when we contrasted averaged signals and cross-location decoding (Fig. S3B).

Temporal dynamics following impulse perturbation was also assessed for impulse 1 and 2 as these preceded cue. At this post-impulse stage, the pattern for memory items was stable across time, as indicated by substantial  temporal generalization (Figure S3C., gray dotted lines indicate decoding above zero, $p < 0.05$). When topographic distribution of memory signal was considered, mostly central posterior electrodes contributed to item classification, regardless of the orientation stimulus having been presented on the left or right, which was again consistent with a spatially generalized code (Figure S4D., Left items, in green; Right items, in pink, $p < 0.05$, one-tailed).

**Figure S3**

*Time-course location specific decoding of Item 1 and Item 2 from voltages, averaged across impulse 1 and 2 during maintenance and associated similarity matrices (SMs) (A), similarity matrices for location-specific and cross-location decoding from impulses (B), temporal generalization of memory representations averaged across Items 1 and 2 relative to impulse 1 and 2 (C); and topographical distributions of memory-related signals for item on the light and left from (100–400 ms) window relative to impulse 1 and 2 onset (D).*

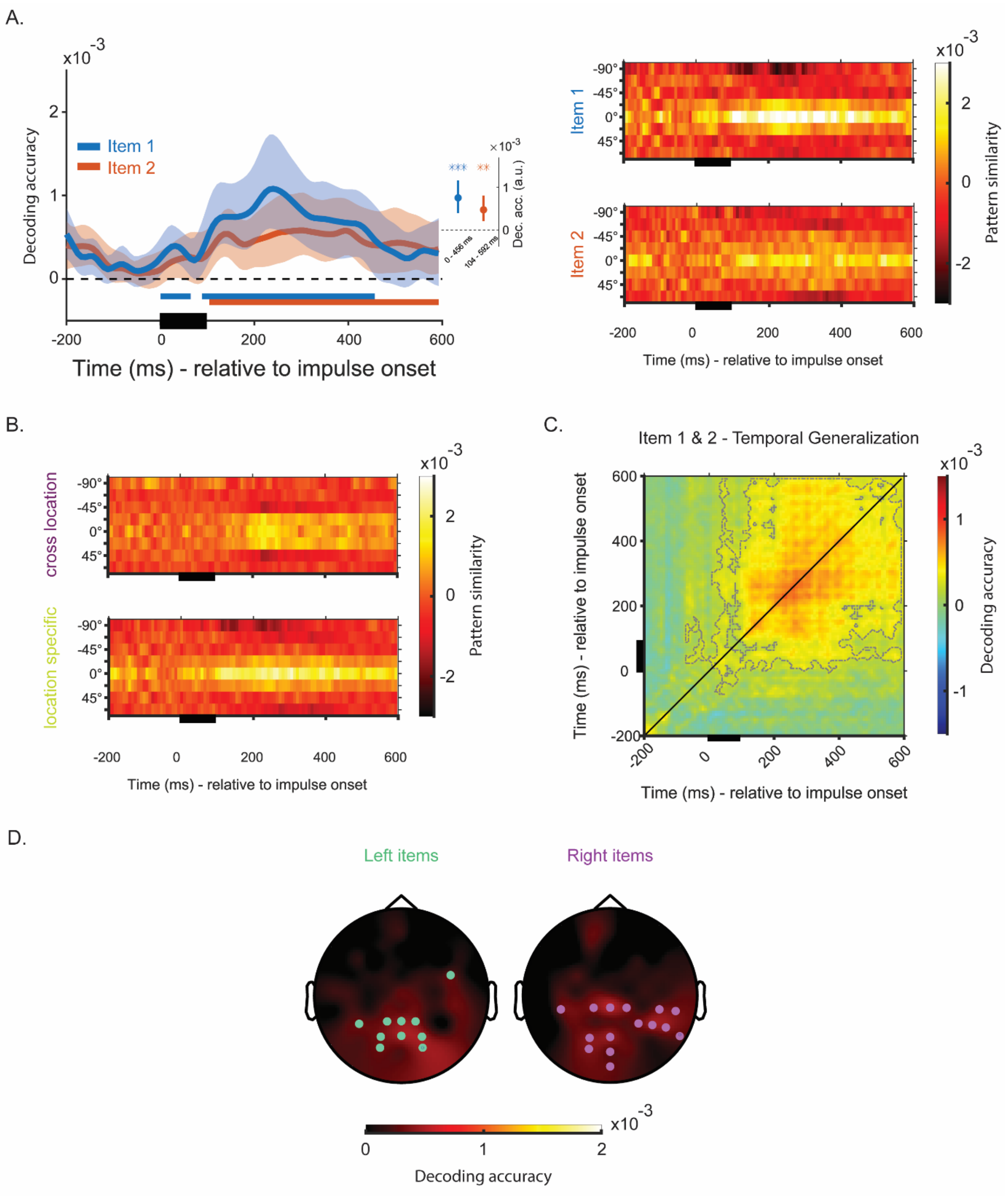

*Note.* **A.** The mean location specific decoding for memory item 1 (dark blue) and Item 2 (orange) from voltages relative to impulse 1 and 2 and associated representational similarity matrices. Colored solid lines represent mean decoding accuracy when training on one spatial location (e.g. left) and testing on other (e.g. right) as a function of time. Shades around solid lines mark 95 % C.I. Solid bars below baseline reflect the time-window with statistically significant decoding according to cluster-based permutation test ($p < 0.05$). Black bars below mark time window with visual stimulation (item or impulse presentation). **B.** Representational similarity matrices for cross location and location specific decoding of voltage data for memory items averaged across item 1 and 2. **C.** Temporal generalization for memory items, averaged across impulse 1 and 2. Areas bordered by dotted lines show the time-zones with above zero decoding with statistical significance ($p < 0.05$). **D.** Topographic distribution of decoding averaged across impulse 1 and 2, revealing electrode contributions for memory items on Left and Right during the initial 100-400 ms window relative to impulses, according to searchlight analysis. Brightness reflects the strength of decoding. Colored dots mark electrodes with above zero decoding accuracy with statistical significance ($p < 0.05$, one-tailed).
